# Bridging physiological responses to population outcomes under variable thermal stress

**DOI:** 10.64898/2026.07.28.741380

**Authors:** Alison J. Robey, David A. Vasseur

## Abstract

Embedding thermal tolerance metrics into population dynamics offers a promising toolkit for understanding the impacts of stressful heat events, but the time-dependence of thermal stress rarely factors into such assessments. Within organisms, heat causes damage that alternatively accumulates under stressful temperatures and is repaired under permissive ones. While risk under exclusively stressful temperatures is well characterized by thermal death time models, resilience to fluctuating temperatures is less clear. Understanding how this damage accumulation within organisms scales up to affect population thermal tolerance is a necessary step for predicting population dynamics and extinction risk. We address this gap by embedding a model of organismal stress and recovery into population dynamics, yielding time-dependent thermal performance curves of population growth rates. By parameterizing this novel framework with existing data from *Drosophila melanogaster*, we explore how integrating thermal stress across biological scales shapes the consequences of organismal stress on population outcomes under realistic thermal fluctuations.

## 1 Introduction

Rising temperatures under anthropogenic climate change are expected to precipitously accelerate species extinction rates (Urban, 2024) as the frequency, intensity, and duration of thermal stress events increase rapidly worldwide (Jørgensen et al., 2022). Temperature’s effects on biological processes are ubiquitous (Kingsolver, 2009), ranging from the rates of molecular processes (Brown et al., 2004) to the carrying capacities of populations (Bernhardt et al., 2018a). Forecasting how novel (and increasingly stressful) thermal regimes will impact organisms, populations, and species thus represents a key goal of current ecology (Buckley et al., 2023; Ellis-Soto et al., 2026). However, realistic integration of thermal stress physiology into metrics of population extinction risk remains a largely unrealized goal, leaving significant uncertainty in our projections of how heat stress may limit population sizes, geographic ranges, and persistence probabilities under climate change.

Comparing measurements of organismal thermal tolerance with projected changes to thermal regimes has been a major focus of recent forecasting efforts. A prevalent approach is predicting the frequency at which organisms experience temperatures exceeding their acute thermal limits (e.g., the critical thermal maximum (*CT*_max_); Sunday et al., 2012; Pinsky et al., 2019) and their likelihood of failure or mortality given the duration and intensity of such exposures (e.g., thermal death time (TDT) models; Rezende et al., 2020; Jørgensen et al., 2022; Willot et al., 2022). Such work characterizes important components of organismal physiology, but its focus on the discrete endpoints of thermal tolerance may not accurately reflect the rate at which thermal damage accumulates over time (Kovacevic et al., 2019; Ørsted et al., 2022), the sub-lethal limitations imposed by sequences of variable temperature stress (Buckley et al., 2022), or the population-level impacts of organismal stress (Clusella-Trullas et al., 2021). Alternative approaches have turned instead to continuous metrics known as thermal performance curves (TPCs), which describe the rise and fall of different rates or traits in response to increasing temperatures (e.g., movement speed, fecundity, fitness; Huey and Stevenson, 1979; Angilletta, 2009). Pairing TPCs with projected frequency distributions (Deutsch et al., 2008; Vasseur et al., 2014, 2025) or sequences (Duffy et al., 2022; Robey and Vasseur, 2026; Robey et al., 2026) of temperatures may provide more nuanced estimates of sub-lethal effects and population-level outcomes like extinction risk.

However, using TPCs for population forecasting relies on several questionable assumptions (Sinclair et al., 2016). First, TPCs are assumed to accurately characterize responses to stressfully hot temperatures, despite typically being poorly parameterized above species’ thermal optima (*T*_opt_; e.g., see the frequently analyzed insect TPC dataset from Frazier et al., 2006). This issue is exacerbated for *population* TPCs because they fundamentally represent equilibrium processes; population growth rate TPCs, for example, must be measured by holding organisms at a constant temperature for at least the duration of their lifespan (thus representing a fully ‘acclimated’ TPC; Huey and Berrigan, 2001; Sinclair et al., 2016). Such TPCs are assumed to translate directly to fluctuating conditions (Bernhardt et al., 2018b), even when those conditions include transient exposures to stressful temperatures (i.e., those temperatures causing population decline) where the rate they aim to capture is inherently unmeasurable at equilibrium (i.e., because the equilibrium population size is zero). Second, TPCs are assumed to depend only on the *current* temperature, despite empirical evidence showing that prior thermal history can greatly affect current thermal performance (‘time-dependent’ or ‘carryover’ effects; Huey and Stevenson, 1979; Somero, 2010; Schulte et al., 2011; Kingsolver et al., 2015; Kingsolver and Woods, 2016; Williams et al., 2016; Kingsolver and Buckley, 2017; Buckley et al., 2025). Depending on the intensity and duration of exposure, stressfully hot temperatures may transiently modify thermal tolerance by inducing beneficial responses, such as acclimation (Kremer et al., 2018; Baeza Icaza et al., 2025) and heat hardening (Loeschcke and Hoffmann, 2007), or harmful ones, such as damage accumulation (Feder and Hofmann, 1999; Tomanek, 2010; Williams et al., 2016; Gonźalez-Tokman et al., 2020). Together, these assumptions suggest that (i) extrapolated TPCs likely underestimate the negative impacts of stressfully hot temperatures and (ii) incorporating thermal history into thermal forecasting may be important for accurately characterizing risk.

Fluctuations in and out of stressful temperatures are ubiquitous in realistic environments, but TPC-based forecasting has historically lacked mechanisms to incorporate physiological plasticity into population dynamics. In contrast, organismal models like TDT frameworks successfully recapitulate damage dynamics under *solely* stressful temperatures, but cannot forecast the organismal outcomes of regimes that fluctuate in and out of stressful temperatures (nor their population-level consequences). Various models have been proposed for bridging this gap: Kingsolver and Woods (2016) incorporated time-dependent effects into growth rate TPCs by modeling the temporal dynamics of stress proteins in the tobacco hornworm *Manduca sexta*, while Klanjscek et al. (2016) created a damage-repair model of cellular-level oxidative damage. More recently, Buckley et al. (2025) modeled thermal damage-repair dynamics under variable temperature fluctuations in the English grain aphid *Sitobion avenae*, while Arnold et al. (2025) proposed a framework for building damage-repair dynamics into TDT models. These frameworks offer critical insights on modeling variable thermal stress, but have generally focused on bridging the effects of sub-organismal processes (e.g., heat shock responses) to organismal-level ones (e.g., larval growth rates, individual survival). We have yet to understand how these temporal dynamics will scale up from organisms to their populations.

Here, we develop a model of how organismal damage-repair dynamics might be incorporated into population-level processes under variable thermal stress. We begin by describing simple damage and repair functions consistent with what is currently known about the temperature dependence of each, then define a population growth rate TPC (comprised of temperature- and damage-dependent birth and death rates) to assess how the short-term dynamics of damage accumulation and repair affect population outcomes over time. We fit our model to previously gathered experimental data from *Drosophila melanogaster* and show that over the permissive range of temperatures (where damage can be maintained at equilibrium), the resulting population TPC is well-defined and follows a classic shape. However, under stressful temperatures (where damage accrual outpaces the capacity for repair), this fully acclimated equilibrium TPC is undefined and population responses depend instead on dynamic processes. This finding represents a major step forward in connecting the thermal physiology observed under stressful conditions (i.e., TDT curves) to classic TPC-based forecasting approaches. We use this conceptually unified model to recapitulate key dynamics under constant conditions and demonstrate how even these simple tradeoffs affect thermal resilience under variable conditions.

## 2 Model Development

Our model is a system of two ordinary differential equations (ODEs) tracking changes to damage level 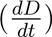 and population size 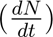 over time *t* in response to changes in temperature *T* (reported throughout in degrees Celsius).

### 2.1 Damage-repair dynamics

Thermal damage arises from mechanisms including protein denaturation, enzyme inactivation, oxygen limitation, osmotic or ionic imbalance, and membrane dysfunction (Schmidt-Nielsen, 1997; Gonźalez-Tokman et al., 2020; Ørsted et al., 2022). In the **stressful temperature range**, damage accumulates rapidly, disrupting homeostasis and eventually causing heat failure or mortality. However, if body temperature returns to the **permissive temperature range**, thermal damage is repaired faster than it accumulates; under permissive temperatures, therefore, heat failure does not occur, allowing for long-term homeostasis and life cycle completion (Ørsted et al., 2022). These temperature ranges are divided by the **critical temperature** *T_c_*: below *T_c_*, temperature limits lifespan indirectly (and gradually) by increasing metabolic and/or aging rates (Shaw and Bercaw, 1962; Maynard Smith, 1963; Brown et al., 2004; Munch and Salinas, 2009), whereas above *T_c_*, temperature limits lifespan directly (and precipitously) by inducing thermal damage (see Fig. S1; Ørsted et al., 2022). We designed a simple model of damage-repair dynamics by employing several key assumptions: first, that all forms of thermal damage are reparable, and second, that accruing or repairing any type of damage results in equivalent energetic costs and gains. Such assumptions may well prove false; for example, heat stress may cause irreversible oxidative damage (Santra et al., 2019) or permanent sterility (Jørgensen et al., 2006), and the diverse suite of physiological stress responses (e.g., synthesis of heat shock proteins, accumulation of reactive oxygen species, DNA repair mechanisms) may be induced at different temperature or damage thresholds with different levels of efficacy and energetic costs (Gonźalez-Tokman et al., 2020; Rennolds and Bely, 2023; Bullard et al., 2026). Even under scenarios where different exposures impose equivalent *amounts* of damage, divergent *mechanisms* of damage may affect the pace and cost of recovery (e.g., short-term exposure to intense stress results in denatured proteins that simply need to be refolded, whereas long-term exposure to moderate stress results in denatured hydrolyzed proteins that need to be replaced altogether; Dingley and Maynard Smith, 1968). Given the complexities and unknowns of this underlying physiology, we use this simple model not to exhaustively characterize organismal damage-repair processes, but rather as a phenomenological description with sufficient detail to recapitulate key dynamics.

When temperatures remain above *T_c_*, heat failure rates have been successfully characterized using TDT (and related thermal tolerance landscape) models. These models assume (i) heat failure rates increase exponentially with temperature (Jacobs, 1919; Fry et al., 1946; Kilgour et al., 1985; Kilgour and McCauley, 1986; Jørgensen et al., 2019b, 2021) and (ii) heat damage is additive (empirically validated in many bacteria species, as well as e.g., brook trout *Salvelinus fontinalis* (Fry et al., 1946), vinegar fly *D. melanogaster* (Jørgensen et al., 2021), and garden thyme *Thymus vulgaris* (Faber et al., 2024)). This additivity (and its insensitivity to exposure order) suggests that the physiological damage mechanisms leading to heat failure are similar under both moderate and severe stress (Ørsted et al., 2022). Given this well-characterized relationship between temperature and damage, we modeled the rate of thermal damage accrual *d* (Fig. 1a) as the sum:

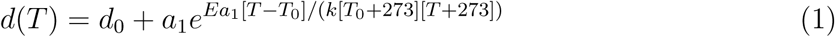

**Figure 1:**
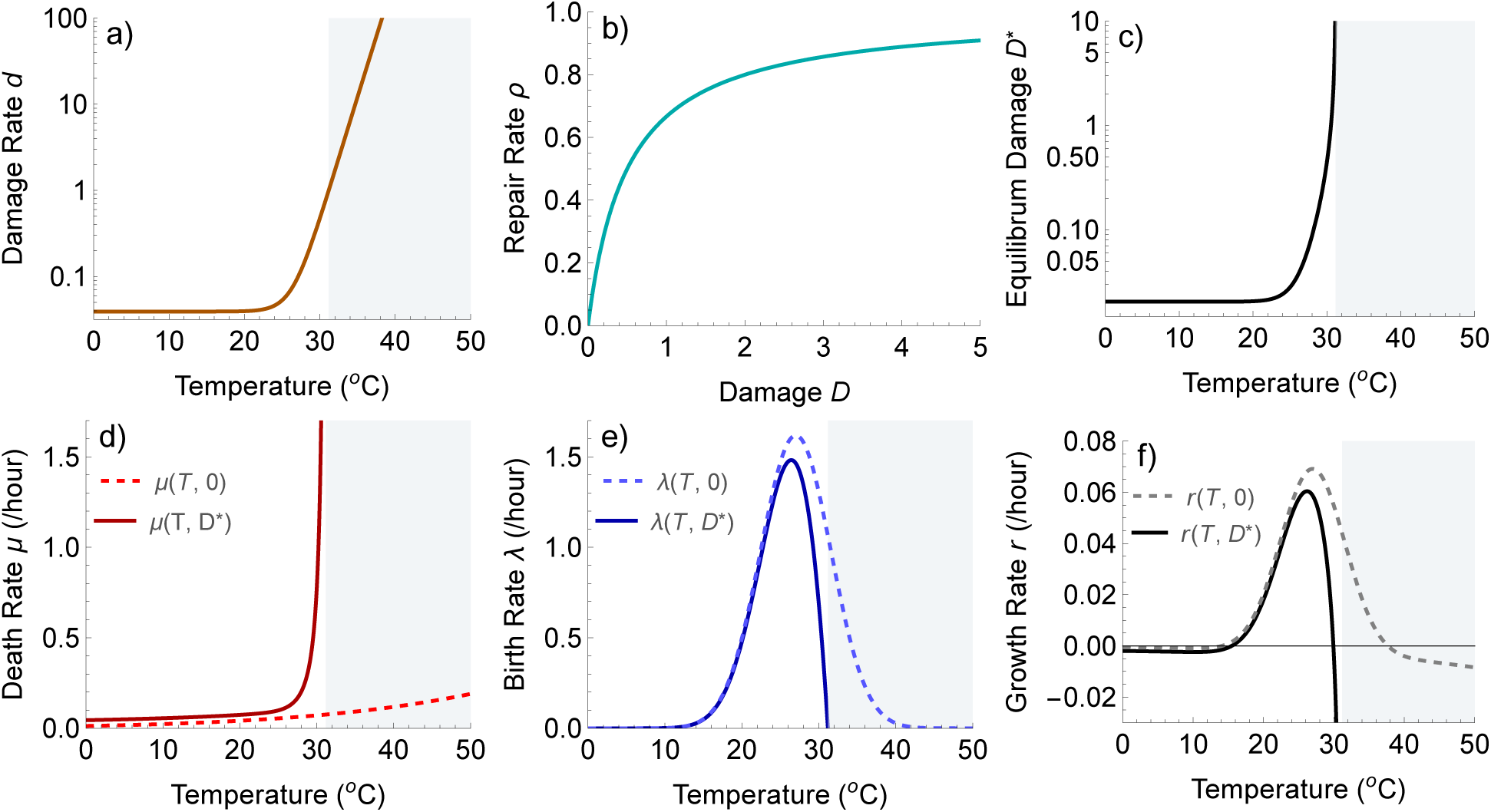
Dynamics of the damage-repair model under constant temperatures (plotted using parameters listed in Table S1). Given the rates of (a) damage at each temperature (Equ. 1) and (b) repair at each damage level (Equ. 2), we solve for (c) the equilibrium amount of damage *D^∗^* at each temperature (Equ. 4). This equilibrium exists for *T < T_c_* (permissive temperatures; white background), but not for *T T_c_* (stressful temperatures; gray background). (d) Death rate (*µ*; Equ. 6), (e) birth rate (*λ*; Equ. 8), and (f) population growth rate (*r*; Equ. 9) in the absence of damage *D* = 0 (dashed lines) versus at equilibrium damage levels *D* = *D^∗^*(*T*) (as plotted in c; solid lines).

**Figure 2:**
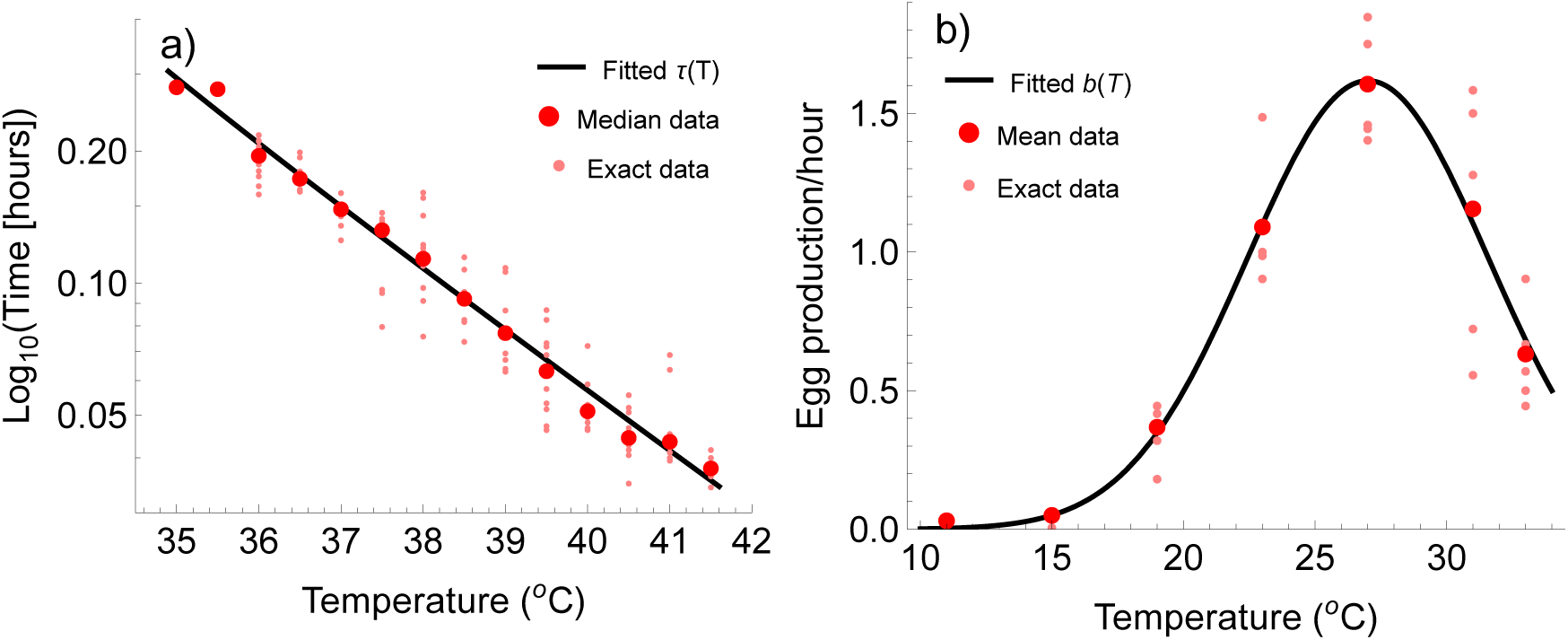
Empirical data (reported median/mean shown in red, underlying data points shown in pink) and fitted functions (black lines) for (a) average death time *τ* (*T*) (Equ. 15; fitted parameters: *d*_0_ = 0.04 damage/hour, *Ea*_1_ = 5.36 eV, *Ea*_2_ = 0.42 eV, *ϕ* = 1.58, *k*_01_ = 3.93 10*^−^*^4^/hour, *k*_02_ = 0.042/hour) compared to median heat failure rates measured across 1-15 individuals per temperature ranging from 35-41.5°(Jørgensen et al., 2019a,b) and (b) birth rate *b*(*T*) (Equ. 7; fitted parameters: *β* = 41.6°, *b*_0_ = 1.62 individuals/hour, *Tb*_opt_ = 27°) compared to average cumulative egg production across 5 individuals per temperature ranging from 11-33°(Overgaard et al., 2014, data extracted via WebPlotDigitizer (Rohatgi, 2024)).

where *d*_0_ represents a low basal rate of damage (e.g., routine oxidative damage) and the second term, a Boltzmann-Arrhenius relationship, represents the temperature-dependent damage rate (where *k* is the Boltzmann constant and *T*_0_ is the intercept temperature). When parameterized by a small intercept *a*_1_ and a high activation energy *Ea*_1_, this term is approximately 0 under most permissive temperatures but accelerates steeply under stressful temperatures, following the expected pattern of sharply accelerating damage rates as temperatures approach and exceed *T_c_* (Ørsted et al., 2022). Given our focus on heat stress, we did not include a mechanism for cold-induced damage here; exploring temperature regimes including cold stress would require a modified U-shaped death function (i.e., instead of one that includes only the ‘right half’ of that U; for more in-depth discussions of cold stress, see Williams et al., 2015; Colinet et al., 2015; Marshall et al., 2020, 2021; Tarapacki et al., 2021; Byrge et al., 2026).

Compared to thermal damage rates, repair rates remain poorly understood and parameterized (Buckley et al., 2025); even the shape of repair functions in response to temperature is uncertain. Previous work has assumed if an organism survived the day, complete repair could occur overnight regardless of nighttime temperature or amount of accumulated damage (e.g., most TDT models; Rezende et al., 2020; Ørsted et al., 2024). Empirical evidence largely disagrees with this assumption, finding instead that both overnight temperature and total stress accumulation alter recovery outcomes (Colinet et al., 2015), for example in blowflies (*Calliphora erythrocephala;* Kashmerry and Bowler, 1977; Bowler and Kashmeery, 1979), English grain aphids (*S. avenae;* Zhao et al., 2014; Ma et al., 2015, 2018), and lady beetles (*Propylea japonica*; Bai et al., 2019). Measurements of actual repair rates are limited, but show some evidence of faster repair at warmer temperatures, including recovery from heat injury in *C. erythrocephala* (Bowler and Kashmeery, 1979) and the bacteria *Staphylococcus aureus* (up to 45°; Iandolo and Ordal, 1966) and *Listeria monocytogenes* (McKellar et al., 1997), as well as recovery from UV damage in *Daphnia pulicaria* (MacFadyen et al., 2004) and *Limnodynastes peronii* tadpoles (Morison et al., 2020). Based on a split-dose experiment across four species of *Drosophila*, Ørsted et al. (2022) suggested repair declines at low temperatures due to slowed metabolism and at high temperatures due to increased disruption rates, a hypothesis supported by recent experiments in *D. suzukii* (Ørsted et al., 2026). Previous models have thus typically assumed accelerating or unimodal relationships between repair rate and temperature (Ørsted et al., 2022; Buckley et al., 2025; Arnold et al., 2025). However, writing repair as a direct function of temperature (i) ignores the biological reality that repair mechanisms need not operate unless damage is present (and, conversely, that repair mechanisms should continue to function under non-damaging temperatures if damage is still present) and (ii) overlooks the potential for time lags between damage accrual and remedy. We thus modeled repair instead as a function of accumulated damage *D* (itself a function of temperature *T*) to more realistically reflect that repair mechanisms (i) need not operate unless damage is present and (ii) will ramp up as the amount of damage increases (e.g., the heat shock response Feder and Hofmann, 1999; Tomanek, 2002). We further assumed that there must be a maximum repair capacity (i.e., a maximum amount of energy available for repair processes) and modeled repair rate *ρ* as a Monod-type function (Fig. 1b):

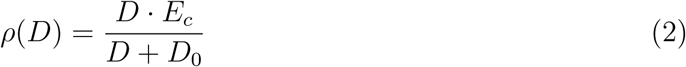

where *D*_0_ is the half-saturation constant and *E_c_* represents proportional available energetic capacity (0 *≤ E_c_ ≤* 1). When damage and repair rates are in equilibrium, repair rate increases with temperature for *T < T_c_*m consistent with empirical evidence (i.e., under constant permissive temperatures, repair rate keeps pace with damage rate as plotted in Fig. 1a); however, repair mechanisms eventually ‘saturate’ for *T > T_c_*, causing the expected reduction in repair capacity given sufficient thermal stress.

Net damage accrual over time is the difference between the temperature-dependent damage rate *d*(*T*) and the damage-dependent repair rate *ρ*(*D*):

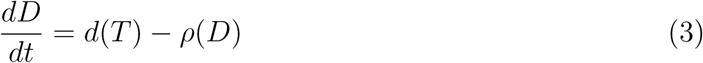

Given a constant temperature *T*, the equilibrium amount of damage *D^∗^* is thus:

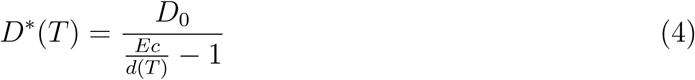

This equilibrium exists for *D^∗^ >* 0 until temperatures reach the critical value where *E_c_* = *d*(*T*) (Fig. 1c); this is the **critical temperature** *T_c_* dividing the permissive range (where repair keeps pace with damage) from the stressful one (where damage is a runaway process). Importantly, *T_c_* here is an output of the model itself, not a parameter estimated beforehand (as in earlier models; e.g., Buckley et al., 2025).

### 2.2 TPCs and population dynamics

Turning to the population-level processes, we modeled population death rate as the sum of metabolic and damage costs. Metabolic costs (*µ_r_*) are well-characterized by a Boltzmann-Arrhenius relationship (as in metabolic theory; Gillooly et al., 2001, 2002; Brown et al., 2004; Savage et al., 2004) with an intercept value *a*_2_ and a low activation energy *Ea*_2_ (such that metabolic costs start higher, but accelerate slower, than damage costs; Fig. 1d):

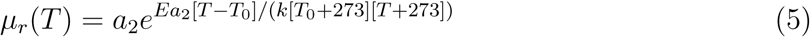

Metabolic costs typically decrease with temperature in the stressful range (Ørsted et al., 2022), so Equ. 5 overestimates the contribution of respiration to death for *T > T_c_*; in this range, however, metabolic costs are negligible relative to damage costs, which we calculate as the amount of damage *D* multiplied by a conversion factor *ϕ*. Since we modeled damage in arbitrary units, *ϕ* represents the rate of death generated by a single ‘unit’ of damage. Overall population death rate thus consists of the additive contributions of metabolic rate and damage, such that death rate increases gradually under permissive temperatures and steeply under stressful ones (as in e.g. Baker and Geider, 2021):

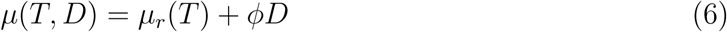

Population birth (or biomass gain) rate (*b*) is typically written as a unimodal function of temperature (Englund et al., 2011; Amarasekare, 2015, 2024; Bieg and Vasseur, 2024). We used a Gaussian function:

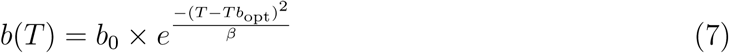

parameterized by an optimal temperature for birth *Tb*_opt_, a breadth parameter *β*, and a conversion factor *b*_0_. Given that the available energetic capacity is *E_c_*, we assumed whatever energy is devoted to repair must be discounted from the overall birth (or biomass gain) rate *λ* (Fig. 1e), such that:

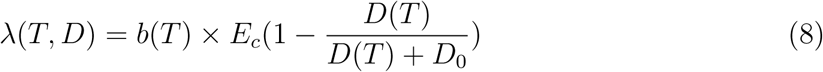

This trade-off between repair and reproduction follows similar patterns observed in experimental systems (e.g., *D. melanogaster* (Krebs and Loeschcke, 1994; Silbermann and Tatar, 2000), *C. elegans* (Aprison and Ruvinsky, 2014), and *Tribolium castaneum* (Sales et al., 2021)) and is consistent with typical energetic models (e.g., dynamic energy budgets): available energy must be allocated between functions, and those functions essential for short-term survival (repair) take precedence over non-essential ones (reproduction). We made the simplifying assumption that *E_c_* is a fixed value divided between only those two functions.

By definition, population growth rate (*r*) is the difference between per-capita birth and death rates (Fig. 1f; as in Amarasekare and Savage, 2012; Thomas et al., 2017; Robey and Vasseur, 2026):

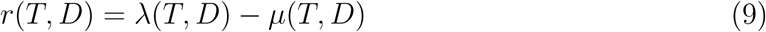

As *µ*(*T, D*) is an increasing function (with the slope dependent on the amount of damage; Fig. 1d) and *λ*(*D, T*) is a hump-shaped function (ranging from symmetric to left-skewed depending on the amount of energy diverted to repair; Fig. 1e), this difference results in a unimodal left-skewed function typical of population growth rate TPCs (Fig. 1f; Rezende and Bozinovic, 2019; Vasseur, 2020). The classical TPC measured for populations is given by this *r*, which here is equivalent to the per-capita growth rate of *N* under constant temperatures prior to the impact of density dependence. The former constraint ensures the TPC does not embody any effects of thermal history, such as physiological or ecological acclimation. This has typically not been an issue in simple models (as acclimation is rarely incorporated, though see Fey et al., 2021), but experiments often assume multiple generations of acclimation are necessary prior to TPC measurement. In our model, exposure to constant temperature is necessary to ensure that damage reaches its equilibrium; the equilibrium TPC is thus given by *r*(*T, D^∗^*(*T*)) (where *D^∗^*(*T*) is the equilibrium damage at a given temperature *T*). Since *D^∗^*(*T*) is undefined above the critical temperature *T_c_*, the TPC is also undefined above *T_c_* (indeed, there is a vertical asymptote at *T* = *T_c_*) because here damage continually increases. Finally, to model temperature-dependent population dynamics, we followed recent precedent (Duffy et al., 2022; Robey et al., 2026; Vasseur et al., 2025; Robey and Vasseur, 2026) in using the temperature-dependent *r*-*α* model of logistic population growth (Long et al., 2007; Mallet, 2012; Vasseur, 2020):

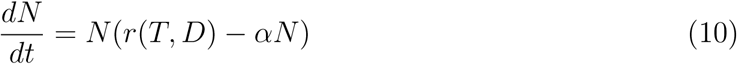

In this model, *α* measures the strength of negative density dependence (which we assume to be independent of temperature, though recent work suggests *α* may increase under generic assumptions; Bieg and Vasseur, 2024) and *r* is the temperature-dependent intrinsic growth rate given by Equ. 9; in contrast to prior work, *r* here depends on the accumulated damage *D* as well as the current temperature *T*. The time-varying equilibrium population size *N^∗^* under permissive temperatures is the temperature-dependent carrying capacity *K*(*T*) = *r*(*T*)*/α* and, under stressful temperatures, is zero.

### 2.3 Model parameterization and comparisons

We explored this model using *D. melanogaster*, for which data are available to estimate most parameters; we used a TDT curve of knockdown times (Jørgensen et al., 2019b,a) to parameterize the death and damage functions (as detailed in Box 1) and a TPC of egg production (Overgaard et al., 2014) to parameterize birth rates (see Supporting Information A, Tables S1, S2). Because this model does not incorporate life stages, development time, or maturation rate, we scaled the population growth rate TPC to an empirically observed maximum rate of *r*_max_ = 0.06/hour (1.45/day; Vinton et al., 2026) to approximate realistic growth. Throughout, we converted time measurements to hours as a compromise between the timescales of thermal damage (seconds to hours), diurnal fluctuations (hours to days), and the organism’s lifespan (weeks to months). While data to estimate the density-dependence parameter *α* is unavailable, the choice of *α* affects only the absolute population size, not the model dynamics (see Supporting Information C, Fig. S3), and we proceeded with *α* = 0.001. Without data to estimate the half-saturation constant of repair rate or the maximum energetic capacity, we set *D*_0_ = 0.5 and *E_c_* = 1 and explored the effects of varying these parameters in section 3.1.

To assess the importance of including organismal damage-repair dynamics in population modeling, we compared model results obtained using the full **dynamic** model (where damage *D* fluctuates throughout simulations and birth-death processes experience the associated trade-offs) to two null models: the **no damage** model (where *D* = 0 throughout simulations, as in the dotted lines in Fig. 1d-f) and the **equilibrium damage** model (where *D* = *D^∗^*(*T*) throughout the simulation, as in the solid lines in Fig. 1d-f). We further compared outcomes to three phenomenological models, where fitness was determined using static TPC model fits (selected from the ‘rTPC’ library; Padfield et al., 2021, 2024) akin to what would be obtained by experimentally measuring populations using typical laboratory protocols: we select five evenly-spaced temperatures at which to ‘measure’ population fitness, sampled 5 times per temperature (assuming normally distributed measurement error), fit each TPC to the sample points, then repeated this protocol 100 times and took the average TPC for each model (see Supporting Information D).

#### BOX 1. Estimating thermal death time

Death rates are often modeled using instantaneous mortality hazard rates *h*(*t*), which are typically assumed to be constant with respect to time, such that *h*(*t*) = *µ*, the mean time to death is *τ* = 1*/µ*, and the survival function (i.e., the probability of surviving a given interval with a known hazard rate) is *S*(*T*) = *e^−µt^* (Ergon et al., 2018; Bullard et al., 2026). However, in this model we have a *time-varying* hazard rate – because morality depends on both current and past conditions via accumulated damage – determined by the death rate function written with respect to time.

Given the choice of a constant temperature *T*_1_, we can thus define our time-varying hazard function:

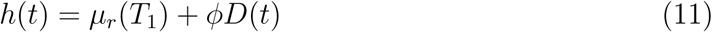

where *µ_r_*(*T*_1_) and *ϕ* are constants dependent on the chosen parameters. Since damage *D* accumulates at a rate 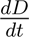 and *D*_0_ is a small constant, as *D → ∞*, *D*_0_ *≪ D* and we can approximate:

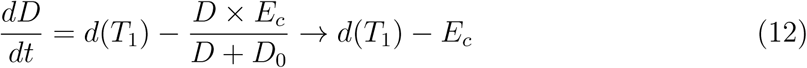

Defining *µ*_0_ = *µ_r_*(*T*_1_) and *µ*_1_ = *ϕ*(*d*(*T*_1_) *− E_c_*), we can write a linear approximation for the time-varying hazard function at a given temperature *T*_1_ as:

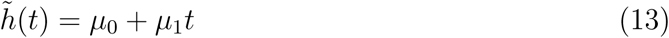

From here, the survival function *S*(*t*) is defined:

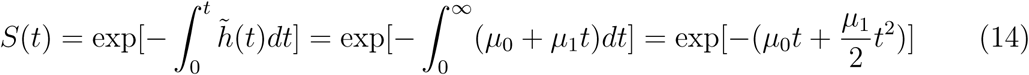

and the average time to death is:

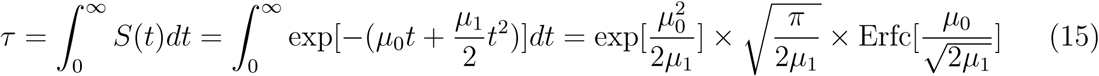

By estimating the average time to death at different constant temperatures *T*_1_ *> T_c_* using Equ. 15, we can generate a TDT curve comparable to commonly measured assays (see Supporting Information B, Fig. S2 for validation of this estimation via Gillespie simulations; Kummel and Vasseur, 2025), which we then fit to experimental data to obtain estimates for all parameters incorporated in Equ. 15.

### 2.4 Simulations

We simulated this system using a numerical solver (*NDSolve* package) in Wolfram Mathematica (V.14.1.0.0). All simulations began at a temperature *T*_0_ *< T_c_* with initial conditions *D*_0_ = *D^∗^*(*T*_0_) and *N*_0_ = *N^∗^*(*T*_0_*, D^∗^*(*T*_0_)). We considered populations extinct at time *t* if *N_t_ ≤* 1 individual. We generated temperature time series composed either of (i) stepwise shifts from *T < T_c_* to *T ≥ T_c_* (to assess persistence time under constant conditions; see section 3.1) or (ii) diurnal fluctuations with various heat stress treatments (to assess resilience under variable conditions; see section 3.2). Non-heatwave diurnal conditions oscillated between ‘nighttime’ temperatures of *T*_min_ *< T* = 20°*< T*_opt_ (hour 0 of each day) and ‘daytime’ temperatures of *T* = 26°*≈ T*_opt_ (hour 12 of each day). We incorporated heatwave treatments of variable length (number of days) and degrees of nighttime and daytime warming (with nighttime temperatures bounded to be at least 1°C cooler than adjacent daytime temperatures) using a smoothing function so that all time series were continuous.

## 3 Model Exploration

### 3.1 Energetic trade-offs and repair capacity

We began by considering the effects of varying the free parameters (energetic capacity *E_c_* and half-saturation constant of repair rate *D*_0_) on model behavior. Reducing *E_c_* and increasing *D*_0_ both increased the equilibrium level of damage *D^∗^* (consequently reducing the equilibrium population size *N^∗^* and the speed of reaching equilibrium), as a lower *E_c_* indicates less energy available for repair whereas a higher *D*_0_ indicates repair mechanisms activating more slowly. Reducing *E_c_* also changed the value of both *T_c_* (the temperature at which damage outpaces repair) and *T*_max_ (the temperature at which death outpaces birth): damage becomes a runaway process at cooler temperatures when energy is limiting compared to when energy is replete. Equilibrium TPCs under energetic limitation had reduced height, breadth, and heat tolerance of *T*_max_ and *T*_opt_ (Fig. 3a), as previously observed under reduced environmental resource levels (e.g., Brett et al., 1969; Brett, 1971; Thomas et al., 2017). Furthermore, given any value of *E_c_*, *T*_max_ *< T_c_* (Fig. 3b), suggesting that individual organisms can survive at hotter constant temperatures than can their overall populations (i.e., there are temperatures *T*_max_ *< T < T_c_* where an individual can survive, but its population cannot persist, in the long-term). The distance between *T*_max_ and *T_c_* increased gradually under greater energy limitation.

**Figure 3:**
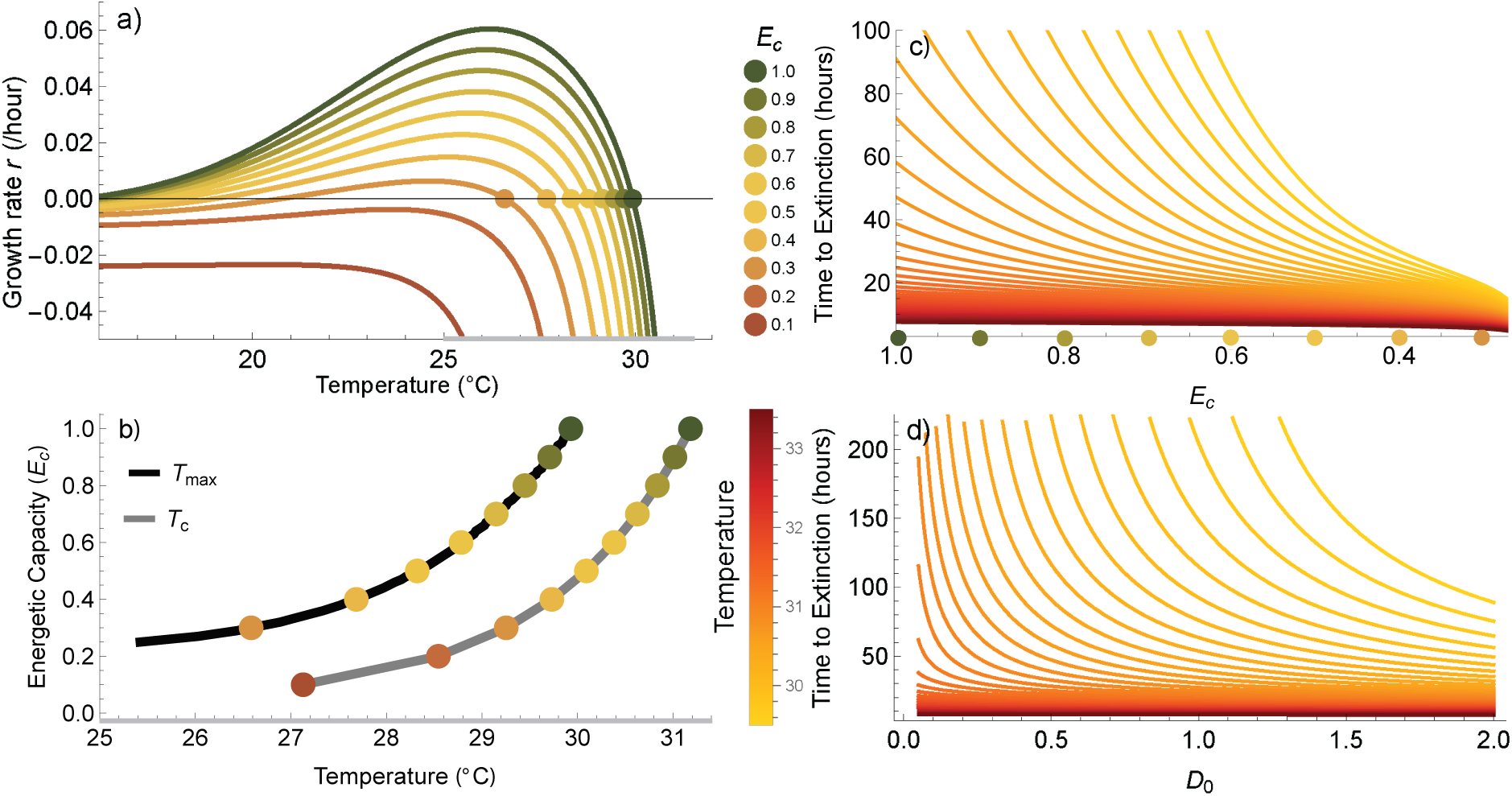
Model sensitivity to varying free parameters. (a) Equilibrium population growth rate TPC under different values of *E_c_*, with *T*_max_ indicated by the color-coded dots. (b) Comparison between *T*_max_ (color-coded dots connected by black line) and *T_c_* (color-coded dots connected by gray line) as *E_c_* is varied; the temperature range plotted across both (a) and (b) is indicated by gray shading on axes. Note that most *T_c_* values would be visible in plot (a) (as each TPC is asymptotic to its corresponding *T_c_*), but are not shown due to overlap between ‘high energy’ *T*_max_ values and ‘low energy’ *T_c_* values. (c, d) Number of hours until extinction after a stepwise shift at *t* = 2 hours (from *T* = 25°to temperatures ranging from 29.5 *≤ T ≤* 33.5°) across different values of *E_c_* and *D*_0_.

Increasing *D*_0_ and reducing *E_c_* both reduced population resilience to step-wise changes in temperatures (Fig. 3c, d). A higher *D*_0_, for example, leads to faster extinctions and extinctions at cooler temperatures compared to a lower *D*_0_ (Fig. 3d). Notably, because we designed the model such that the energetic capacity is divided between repair and reproduction, very low values of *D*_0_ (i.e., very fast reallocation of energy from reproduction to repair) can lead to transient reductions in birth rates and abundance prior to reaching equilibrium.

### 3.2 Effects of time-dependent thermal tolerance on population dynamics

Predictably, under diurnal temperature fluctuations with variable heat stress treatments, the dynamic damage model was less resilient to heat stress than the no damage model and more resilient than the equilibrium damage model (comparison between teal, black, and gray lines in Fig. 4a-c). Extinction risk was most sensitive to the extent of daytime warming, but whether a given amount of daytime warming caused extinction depended on both the heatwave duration (e.g., dynamic model can survive through one, but not two, days of the heatwave treatment in Fig. 4a) and the extent of nighttime warming (gradients across Fig. 4e). If daytime warming was not significant (e.g., *<* +4.8°, equivalent to peak temperatures of 30.8°) or too significant (e.g., *>* +6.7°, equivalent to peak temperatures of 32.7°), extinction was no longer sensitive to heatwave duration or nighttime warming: in the former case, temperatures never surpass *T_c_* and in the latter case, they remain above *T_c_* for too long to permit organism survival. For these models, no duration effects were observed beyond a heatwave length of three days.

**Figure 4:**
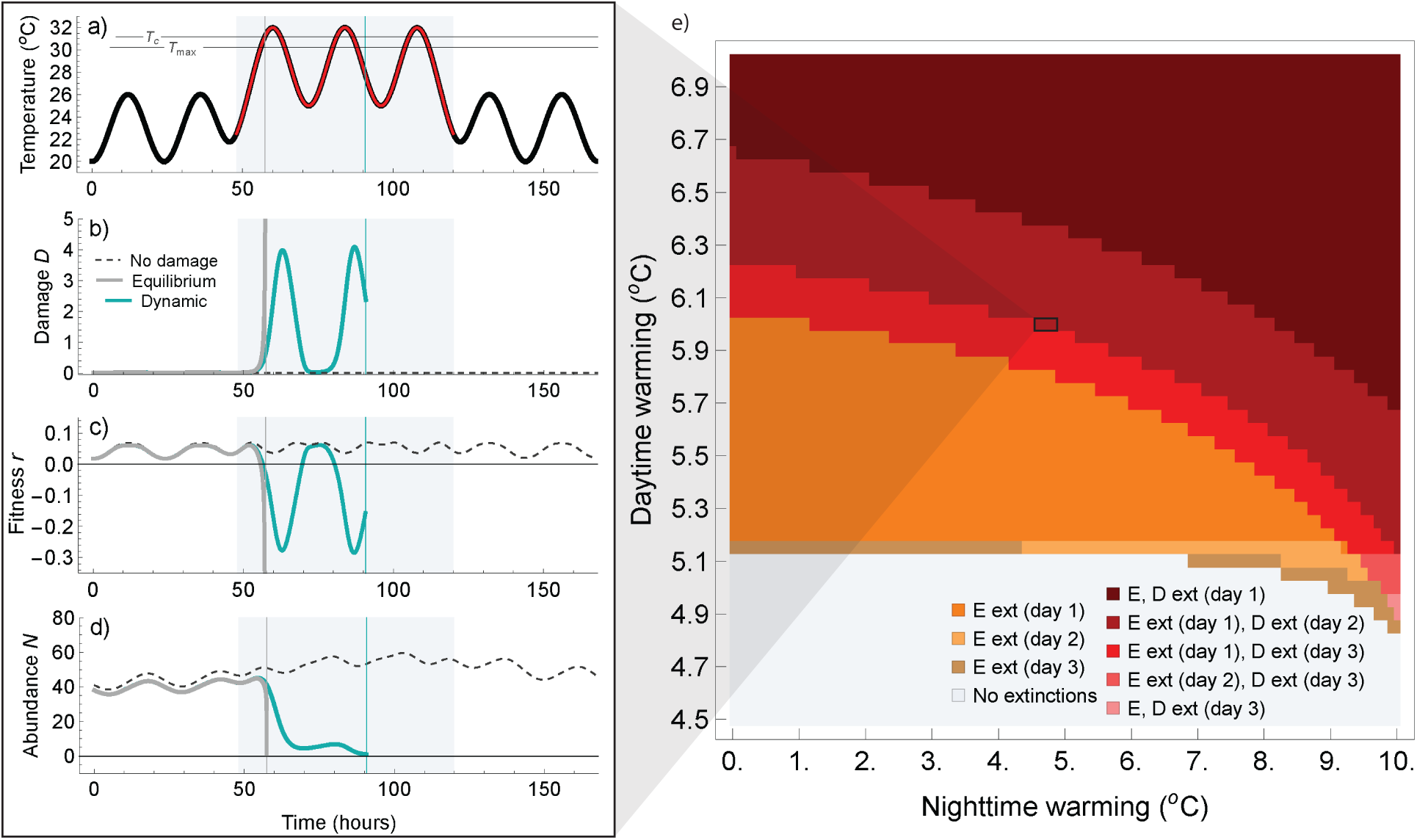
Comparison of dynamic model with null models across heatwave treatments. (a) Example temperature time series with a 3-day heatwave (red; daytime +6°, nighttime +5°) starting on day three (indicated throughout by shaded area); *T*_max_ and *T_c_* are indicated by horizontal lines. The resulting (b) damage, (c) fitness, and (d) abundance over time under the **no damage** model (dashed), the **equilibrium damage** model (gray), and the **dynamic damage** model (teal) with simulations plotted up until extinction (vertical lines). (e) Extinction outcomes under each model given different combinations of nighttime and daytime warming. Both the **equilibrium** (E) and **dynamic** (D) models went extinct in red regions, while only the **equilibrium** (E) model went extinct in orange regions (no extinctions were recorded for the **no damage** model); shading indicates the day on which extinction occurred.

We further compared projected resilience under the dynamic model to three static phenomenological models: (1) the double exponential **Lactin2** model (Lactin et al., 1995), which underestimated fitness slightly above *T*_max_; (2) the exponential-quadratic **Thomas** model (Norberg, 2004; Thomas et al., 2012), which accurately estimated fitness above *T*_max_; and (3) the Gaussian-quadratic **Deutsch** model (Deutsch et al., 2008), which marginally overestimated fitness above *T*_max_ (Fig. 5a). Across the various warming treatments (e.g., Fig. 5b), we found that extinction risk was predictably sensitive to mis-characterizations of fitness above *T*_max_: the TPC with the highest fitness under hot temperatures (Deutsch model; yellow) projected the most resilience to heat stress, followed by the Thomas model (red), the dynamic model (teal), and the Lactin2 model (purple; Fig. 5a). Despite the Thomas model appearing to characterize heat stress well, it routinely overestimated population resilience to daytime warming by around 2°, while the Deutsch model (which overestimated *T*_max_ by only 0.4°) overestimated resilience significantly (Fig. 5c, red and yellow compared to teal). While the Lactin2 model underestimated *T*_max_ slightly, it projected the resilience of the dynamic model very closely (purple compared to teal).

**Figure 5:**
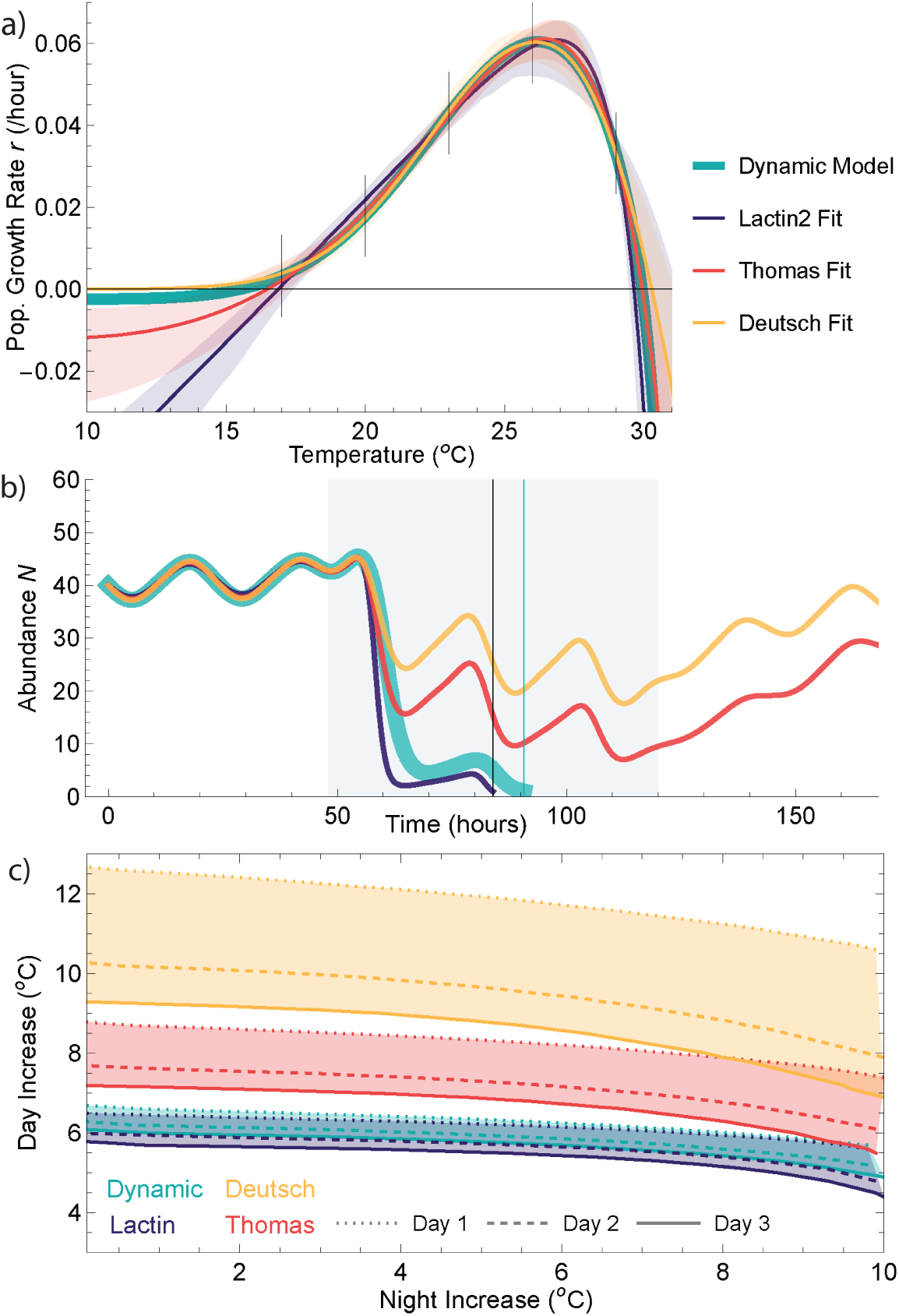
Model outcomes compared to different TPC model fits. (a) Comparison between the ‘true’ TPC (given by the dynamic damage model, teal; *T*_max_ = 29.9°) compared to three phenomenological TPC fits (Lactin2 in purple, *T*_max_ = 29.6°; Thomas in red, *T*_max_ = 29.9°; and Deutsch in yellow, *T*_max_ = 30.3°), each the average of 100 different fits to five replicate measurements at each test temperature (indicated by vertical lines, plotted from 2 standard deviations to the true value), shown with 95% confidence intervals (shading). (b) Comparison between population dynamics modeled using each different TPC fit given a three-day heatwave (daytime +6°, nighttime +5°; shown in Fig. 4a). (c) Projected extinction boundaries under each model given different combinations of nighttime and daytime warming, with day of extinction indicated by line type. Given a TPC model and the combination of day-time and nighttime warming, populations below all corresponding boundaries are expected to persist, those between corresponding boundaries are expected to experience duration effects (i.e., survive one or two days before extinction), and those above all corresponding boundaries are expected to go extinct on the heatwave’s first day.

## 4 Discussion

We have developed a simple, conceptually unified mathematical framework for linking thermal tolerance across biological scales (organism to population) and thermal regimes (permissive to stressful). This model connects population growth rate TPCs measured under permissive temperatures to TDT curves under stressful ones, and offers a straightforward interpretation of the constant temperatures *T*_max_ and *T_c_* at which chronic population persistence and individual survival, respectively, fail. By linking organismal damage-repair processes to population birth-death processes, we show that the time-dependence of organismal thermal tolerance scales up to impact the resilience of their population to thermal stress under dynamic temperature fluctuations.

This framework recapitulates several key hypotheses about resilience to heat stress. First, we find that individual organisms consistently survive at hotter constant temperatures than their populations can persist at (i.e., *T*_max_ *< T_c_*). This follows the hypothesis that thermal tolerance measured at higher levels of biological organization (i.e., populations) should be narrower and less heat tolerant than those measured at lower levels of biological organization (i.e., organisms; Dell et al., 2011; Rezende and Bozinovic, 2019; Bozinovic et al., 2020), an observation further consistent with patterns of ‘thermal tolerance limits’ exceeding ‘thermal fertility limits’ in the TDT literature (Walsh et al., 2019; Ørsted et al., 2024). This difference is an important reminder that forecasting population outcomes (e.g., extinction or extirpation risk) using sub-population level TPCs – a common practice, as most available TPCs measure organismal or sub-organismal rates (Sinclair et al., 2016; Kontopoulos et al., 2024; Cicchino et al., 2026) – will likely overestimate the amount of heat stress a viable population can withstand. This effect could potentially be tempered, however, by intraspecific variation in thermal tolerance: such variation could elevate *T*_max_ if more heat tolerant phenotypes contribute disproportionally to population growth. Careful consideration of what is measured by a given thermal tolerance metric, how closely that trait is linked to long-term population viability, and how it may be affected by realistic sequences of thermal stress are important factors to consider when extrapolating from available thermal tolerance data.

Second, we find that population TPCs show reduced height, breadth, and thermal maxima under energetic limitation, a pattern that has frequently been noted under reduced environmental resource levels (e.g., Brett et al., 1969; Brett, 1971; Thomas et al., 2017; Rueda Moreno and Sasaki, 2023). As pointed out by Huey and Kingsolver (2019), under these constraints, an individual with restricted food intake will also have restricted thermal tolerance, reducing the amount time (and potentially space) where temperatures permit foraging activities, leading to further restriction of food availability; if this positive feedback cycle perpetuates, ectotherms face “metabolic meltdown.” In our model, this pattern reemerges at the population scale. Because we have assumed a fixed energetic capacity *E_c_* within organisms (where survival (via repair) is prioritized over reproduction), under food limitation (reduced *E_c_*), damage outpaces repair under cooler temperatures within organisms (*T_c_* shifts left) and death outpaces birth under cooler temperatures within populations (*T*_max_ shifts left). The repercussions of increasing thermal stress may thus be greatly amplified by food limitation for both organisms (Huey and Kingsolver, 2019) and their populations. Understanding how such patterns could affect consumer-resource dynamics (Vinton and Vasseur, 2022), food web dynamics (Gibert et al., 2022), and community responses (Kremer et al., 2025) will be important next steps in projecting ecosystem responses to increasing heat stress.

Third, we find that population resilience to thermal stress under diurnal temperature fluctuations depends on recovery temperature (nighttime warming) and stress duration (heat-wave length), as well as on peak daytime warming. Earlier models including the sequence, duration, and intensity of thermal stress (e.g., Ma et al., 2015; Buckley et al., 2025) predicted such effects, but such dependence had yet to be shown in a population model with mechanisms for incorporating thermal damage and repair; this result reinforces the conclusion that forecasting risk using only thermal distributions (e.g., Deutsch et al., 2008; Vasseur et al., 2014; Estay et al., 2014; Slein et al., 2023) may underestimate the potential impacts of intensifying heatwave regimes (Robey and Vasseur, 2026; Robey et al., 2026). Incorporating damage and repair processes into such temporally explicit modeling also indicates the importance of better understanding the thermal dependence of repair processes (Arnold et al., 2025; Buckley et al., 2025; Ørsted et al., 2026): warming permissive temperatures may be either beneficial (e.g., if warming brings temperatures closer to *T*_opt_, speeding population growth and repair rates) or harmful (e.g., if higher metabolic costs exhaust energetic resources even the absence of resource decline, such as observed over seasonal cycles in diapausing insects (Williams et al., 2016)). The negative impacts of nighttime warming projected by our model indicate the importance of considering temperature fluctuations over time (and their effects on energy budgets) wholistically. While capturing fully detailed responses to such cycles required a model incorporating damage accrual, repair, and ensuing demographic tradeoffs, we do find that phenomenologically fit TPCs – when they accurately characterize realized fitness at and above *T*_max_ – approximate extinction outcomes reasonably well, suggesting that well-parameterized TPCs may still provide exceptionally useful forecasting tools.

While this model is a useful step forward in our ability to link organismal to population-level processes under variable thermal stress, several key limitations and possible extensions should be considered for further use. Many simplifying assumptions were made in the model setup, particularly in the scaling of organismal damage and repair processes into energy usage and subsequent demographic processes (discussed in detail in Model Development); more data on rates of temperature-dependent damage accrual and repair is needed to generate more nuanced and testable models, but such data is notably difficult to collect (Ørsted et al. (2022), though see Ørsted et al. (2026)). Previously introduced models handling the complex physiological responses to thermal stress with more detail (e.g., Kingsolver and Woods, 2016) are better suited to questions probing the dynamics of within-organism processes. Furthermore, our focus here has been exclusively on the *negative* consequences of time-dependent thermal tolerance; understanding the potential *positive* consequences – namely, beneficial thermal acclimation – remains to be carefully considered, but may be an extremely important facet of realized thermal risk under realistic temperature fluctuations. Explicit incorporation of the time-dependence of phenomena including the heat shock response, heat hardening, and rapid acclimation will be insightful next steps.

Our model also includes simplifying assumptions about how thermal tolerance scales from individuals to populations; namely, we have assumed all individuals in our population (i) have identical thermal tolerance and physiological responses and (ii) are uniformly (and simultaneously) exposed to the exact same temperature sequences. Intraspecific variation in thermal tolerance is, however, widely documented (e.g., Blanchard et al., 2024; Cocciardi and Ohmer, 2024) and, depending on the structure of that variation, may have diverse consequences for the thermal tolerance of the overall population (Lu et al., 2025). Furthermore, individuals are typically dispersed across spatially heterogeneous landscapes and may therefore have differential inherent exposures to stressful temperatures (based on geography, topography, etc.) or abilities to avoid unfavorable temperatures through behavioral thermoregulation (Huey and Slatkin, 1976; Sears et al., 2011, 2016). Both intraspecific variation and the spatial distribution of thermal resources may affect the evolution of thermal tolerance over time: exposure to extreme heat events, even at low frequencies, is known to substantially affect evolutionary trajectories (Buckley and Huey, 2016; Williams et al., 2016), but – particularly for motile organisms in thermally heterogeneous landscapes – avoiding such extremes through behavioral thermoregulation may greatly reduce the pace of heat tolerance evolution (i.e., the Bogert effect; Bogert, 1949; Huey et al., 2003; Muñoz, 2022). Populations’ potential to handle climatic changes at longer timescales will depend heavily on whether (and how quickly) thermal tolerances evolve to match with novel environmental conditions (Vinton and Vasseur, 2020; Urban et al., 2026). The potential for rapid evolution of thermal tolerance (Malusare et al., 2023), however – particularly when species have the requisite ‘tools’ at their disposal (i.e., high spatial thermal heterogeneity through connected, preserved habitats (e.g., Fey et al., 2019); high standing genetic variation through large population sizes (e.g., Forester et al., 2022)) – highlights the critical importance of conservation efforts aimed at supporting large, viable populations to give species their best chance at surviving changing thermal regimes. Better understanding the time-dependence of thermal tolerance is crucial to forecasting such dynamics.

Here, we have established a novel framework for linking the time-dependence of dynamic organismal and population-level thermal responses. Current forecasting approaches largely rely on extrapolating population TPCs (generally with poorly-parameterized decline rates) into stressful temperature regimes where equilibrium population sizes and decline rates fundamentally do not exist. Frameworks that allow us to better predict the effects of heat stress by leaning on different types of data – here, physiological assays of heat failure rates – offer a promising approach to fill this knowledge gap. Furthering such frameworks to incorporate the realistic variation and timescales of thermal stress will be essential to better forecasting the resilience of organisms, populations, species, and communities to changing climates.

## Data Accessibility

All code used in this study are available in Zenodo at doi.org/10.5281/zenodo.21540084

## Acknowledgments

Funding for this study was provided by Yale University. We thank Martina Dal Bello, Colin Kremer, and Martha Muñoz for their helpful feedback on earlier versions of this manuscript.

