## Supporting Information for "Bridging physiological responses to population outcomes under variable thermal stress"

July 29, 2026

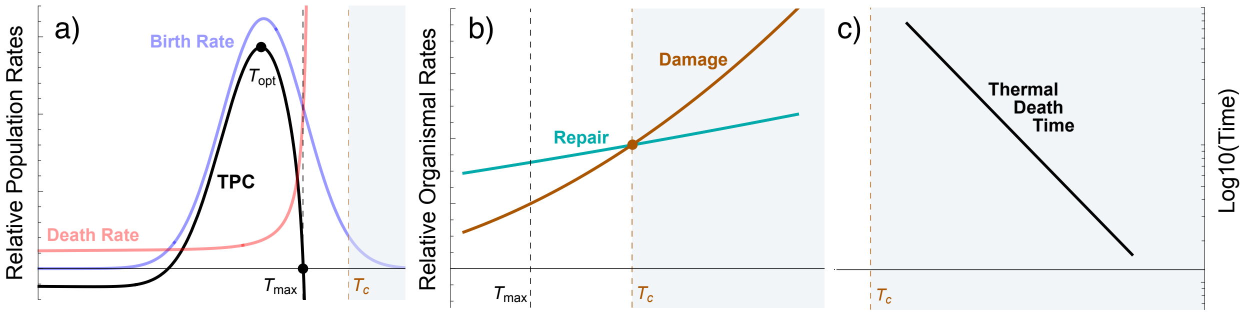


Figure S1: Illustration of various thermal tolerance metrics under different thermal regimes. (a) Under permissive temperatures (white background), tolerance is measured via equilibrium thermal performance curves (TPCs) of population growth rate (equivalent to birth rate less death rate), which peak at the thermal optimum ($T_{\text{opt}}$) and have a right intercept ($T_{\max}$) representing the constant temperature at which the equilibrium population size transitions from a positive value (persistence) to 0 (extinction). (b) Permissive temperatures (white background) are divided from stressful temperatures (gray background) by the critical temperature ($T_c$), where homeostasis is lost when damage (or disruption) rate outpaces repair (or homeostatic capacity) rate (recreated from Ørsted et al., 2022). This is the constant temperature at which the equilibrium individual outcome transitions from survival to death. (c) Above $T_c$, organisms are in the stressful temperature range and governed by fundamentally non-equilibrium processes. Thermal tolerance is measured using thermal death time (TDT) curves describing the exponential relationship between temperature and time to failure (frequently shown as a linear regression via a $\log_{10}$ transformation).

### A Fitting parameters and protocol

| variable | definition | units |
|---|---|---|
| $b$ | maximum per capita birth rate | /unit time |
| $d$ | damage rate | [arb. unit]/unit time |
| $D$ | amount of damage | [arb. unit] |
| $D^*$ | equilibrium damage level | [arb. unit] |
| $\lambda$ | per capita birth rate | /unit time |
| $\mu$ | per capita death rate | /unit time |
| $\mu_r$ | metabolic rate costs | /unit time |
| $N$ | population size | individuals |
| $r$ | per capita population growth rate | /unit time |
| $\rho$ | repair rate | [arb. unit]/time unit |
| $t$ | time | time unit |
| $T$ | temperature | °C |
| **parameter** | **definition** | **value** |
| $\alpha$ | intraspecific density dependence | 0.001 |
| $a_1$ | intercept value of damage rate | **3.93×10$^{-4}$/hour** |
| $a_2$ | intercept value of metabolic rate | **0.042/hour** |
| $\beta$ | birth function breadth parameter | **41.62°C** |
| $b_0$ | birth rate conversion factor | **1.62 individuals/hour** |
| $d_0$ | basal damage rate | **0.04 [arb. unit]/hour** |
| $D_0$ | repair rate half-saturation constant | 0.5 |
| $Ea_1$ | activation energy of damage rate | **5.36 eV** |
| $Ea_2$ | activation energy of metabolic rate | **0.41 eV** |
| $E_c$ | maximum energetic capacity | 1 |
| $\phi$ | conversion factor | **1.58/([arb. unit]×hour)** |
| $k$ | Boltzmann Constant | $8.6173 \times 10^{-5}$ eV/K |
| $r_{\text{max}}$ | maximum height of fitness TPC | 1.45/day |
| $T_0$ | rescaled intercept temperature of BA rates | 20°C |
| $Tb_{\text{opt}}$ | optimal reproduction temperature | **27°C** |
| $T_c$ | critical temperature | **31.18°C** |

Table S1: List of all variables and parameters, their definitions, and (if applicable) their set values and/or units (fitted parameters shown in bold).

Fitting parameters of functions to data in *Mathematica* via *NMinimize* requires setting upper and lower boundaries on the allowable outcomes of each parameter. As the chosen boundaries can significantly affect the fit, we list the exact boundaries used in Table S2. Fits were rejected if (1) *NMinimize* returned "Failed to converge to the requested accuracy or precision within 100 iterations," indicating no reasonable fits within the provided parameter boundaries; (2) fits were calculated, but chosen parameter values were equivalent to either the minimum or maximum value available, indicating that the optimal value may be outside the provided parameter range; (3) visual comparison between fits and datasets was poor; or (4) the resulting $T_c$ fell outside the expected region of 30-35°C.

| **parameter** | **fitted value** | **fit boundaries** | **fitted to** |
|---|---|---|---|
| $\beta$ | 41.6189°C | [0, 500] | $b(T)$ |
| $b_0$ | 1.61789 individuals/hour | [0, 200] | $b(T)$ |
| $d_0$ | 0.039178 damage/hour | [$10^{-4}$, 0.1] | $\tau(T)$ |
| $Ea_1$ | 5.35722 eV | [4.42, 8.82] | $\tau(T)$ |
| $Ea_2$ | 0.413638 eV | [0.28, 0.71] | $\tau(T)$ |
| $\phi$ | 1.57845 damage/hour | [$5 \times 10^{-5}$, 5] | $\tau(T)$ |
| $k_{01}$ | $3.93871 \times 10^{-4}$/hour | [$6 \times 10^{-5}$, $6 \times 10^{-3}$] | $\tau(T)$ |
| $k_{02}$ | 0.04146/hour | [$6 \times 10^{-3}$, 6] | $\tau(T)$ |
| $Tb_{\text{opt}}$ | 27.0093°C | [20, 30] | $b(T)$ |

Table S2: List of fitted parameters, their exact fitted values, the limits placed on possible value choices, and the function they were fitted to. Parameters fitted to $b(T)$ and $\tau(T)$ used data from Overgaard et al. (2014) or Jørgensen et al. (2019a,b), respectively.

### B TDT Validation

Fitting the $\tau(T)$ function required us to generate a thermal death time (TDT) curve from our ODE system using a linear approximation of the time-varying hazard rate $h(t)$ and subsequent approximations of the survival function $S(t)$ and average time to death $\tau(T)$ (given a constant temperature $T > T_c$), as detailed in main text Box 1. To validate this method, we first tested that the linear approximation $\tilde{h}(t) = \mu_0 + \mu_1 t$ (main text Equ. 13) matched the outputs of our used death rate function $\mu(T, D)$ well for $T > T_c$ (Fig. S2a). We then compared the average time to death predicted using $\tau(T)$ versus the average time to death observed across 1000 Gillespie simulations per temperature (Fig. S2b). The approximation matched realized dynamics well, particularly under more stressful temperatures. At temperatures closer to $T_c$, the linear approximation was (predictably) less precise because the total amount of damage $D$ is relatively smaller, leading the approximation of $\frac{D \times E_c}{D + D_0} \to E_c$ (and the subsequent calculation of $\mu_1$) to be a worse approximation of the system's behavior.

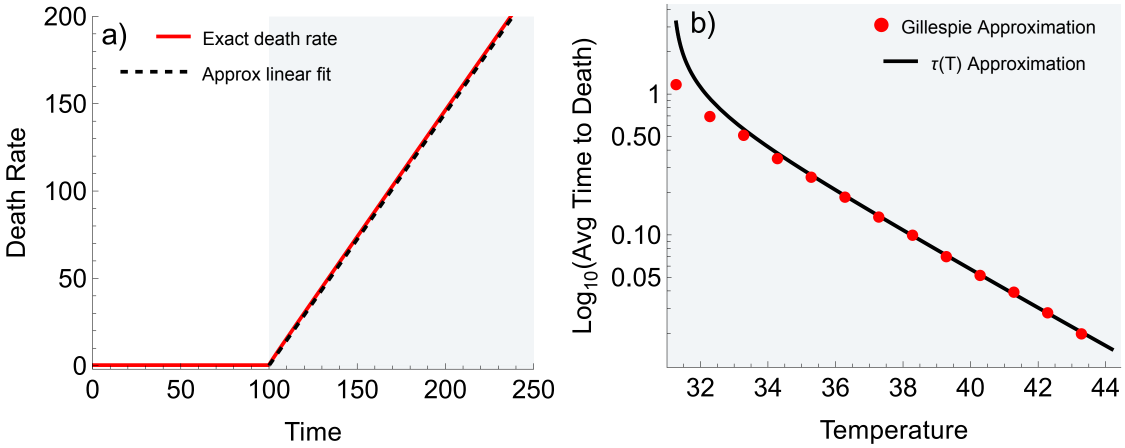


Figure S2: (a) Given a shift from $T = T_c - 1$ to $T = T_c + 1$ at $t = 100$, comparison between the exact death rate ($\mu(D, T)$, main text Equ. 6) modeled via ODE simulations (red) and the linear approximation of the time-varying hazard function (black, main text Equ. 13). (b) Example comparison between the average time to death calculated using Gillespie simulations of the ODE system (red) versus the approximated time to death function $\tau(T)$ (black, main text Equ. 15).

### C Insensitivity to variation in $\alpha$

As we did not have data available to estimate the value of the density-dependence parameter $\alpha$ for the $r$-$\alpha$ formulation of the logistic growth model, we calculated model results throughout using $\alpha = 0.001$. As shown in Fig. S3, changing the value of $\alpha$ affects the magnitude, but not the behavior, of population dynamical outcomes. Differences in extinction timing are generated if the extinction boundary used is not changed in tandem with the choice of $\alpha$ (e.g., in the main text, we considered populations extinct at time $t$ if $N_t \leq 1$, equivalent to $1/\alpha \times 10^{-3}$; extinction outputs remain unchanged with respect to $\alpha$ if this boundary is held at $1/\alpha \times 10^{-3}$, but vary if this boundary is held arbitrarily at 1).

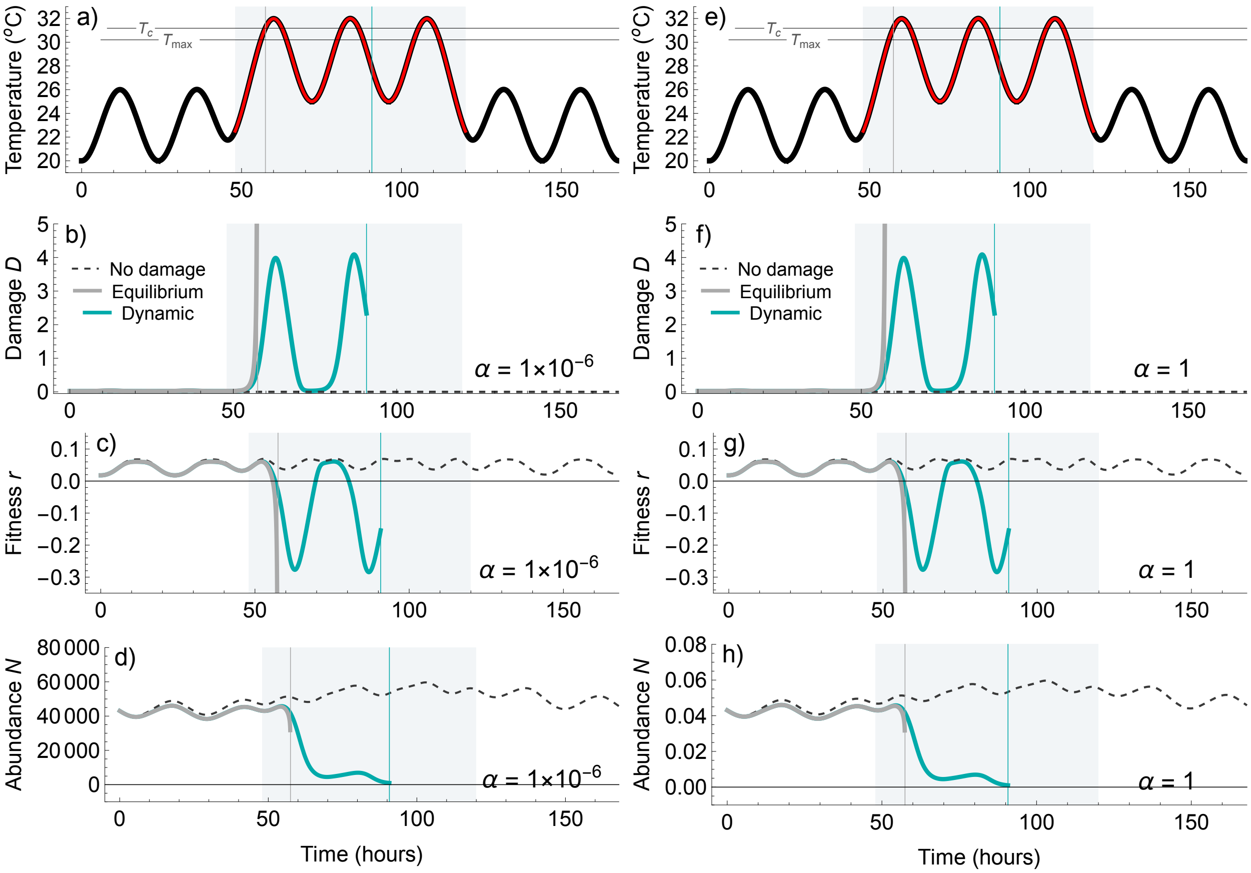


Figure S3: Example of the insensitivity of model results to a six-degree magnitude change of $\alpha$, with dynamics under (a-d) $\alpha = 1 \times 10^{-6}$ on the left and (e-h) $\alpha = 1$ on the right under the same temperature scenario as Fig. 5. (b, f) Damage and (c, g) fitness dynamics are identical, while (d, h) abundance dynamics have a different magnitude but equivalent behavior. Populations are considered extinct at time $t$ if $N_t \leq 1/\alpha \times 10^{-3}$

.

### D Phenomenological TPCs

Assuming the dynamic TPC model (Equ. 9; Fig. 1f, solid line) obtained using the full set of damage-repair and birth-death functions is the 'true' fitness TPC of our example species, experimentally measuring that TPC would typically involve fitting a phenomenological TPC model formulation to replicate measurements of fitness at several different temperatures. To assess whether such protocols might accurately characterize the dynamics shown here, we selected five evenly-spaced temperatures at which to 'measure' the population growth rate: three between $T_{\min}$ and $T_{\text{opt}}$ ($17, 20, 23$°C), one approximately equal to $T_{\text{opt}}$ (26°C), and one between $T_{\text{opt}}$ and $T_{\max}$ (29°C), consistent with frequently used protocols (see e.g., datasets compiled in Frazier et al., 2006; Kontopoulos et al., 2024). We then 'sampled' growth rates 5 times per temperature (assuming measurement error to be normally distributed around the 'true' population growth rate given by Equ. 9). We repeated this protocol 100 times and fitted TPCs to each set of sample points, then took the average of those fits as the 'measured' TPC.

From the 'rTPC' library (Padfield et al., 2021, 2024), we selected three frequently used unimodal TPC formulations which fit the data well (and, crucially for population dynamics, which allowed negative values outside of the thermal breadth):

- The **Lactin2** model is defined:

$$r_L(T) = \exp[aT] - \exp[aT_{\max} - \frac{T_{\max} - T}{\delta_t}] + b \tag{1}$$

  where $a$ determines the steepness of the curve's rise, $b$ determines its height, $T_{\max}$ is the temperature $> T_{\text{opt}}$ at which the curve decelerates, and $\delta_t$ is the thermal safety margin (Lactin et al., 1995).

- The **Thomas** model is defined:

$$r_T(T) = ae^{bT}(1 - (\frac{T - T_{ref}}{c/2})^2) \tag{2}$$

  where $a$ and $b$ are arbitrary constants, $c$ is the thermal breadth, and $T_{\text{ref}}$ dictates the maximum of the quadratic term (Thomas et al., 2012).

- The **Deutsch** model is defined:

$$r_D(T) = \begin{cases} r_{\max} \exp[-\frac{T-T_{\text{opt}}}{2a}^2], T < T_{\text{opt}} \\ r_{\max}(1 - (\frac{T-T_{\text{opt}}}{T_{\text{opt}}-T_{\max}})^2), T > T_{\text{opt}} \end{cases} \tag{3}$$

  where $r_{\max}$ is the maximum fitness at the optimum temperature $T_{\text{opt}}$, $T_{\max}$ is the right intercept, and $a$ determines the curve breadth (Deutsch et al., 2008).

Average curve fits were insensitive to increasing the number of sample points (beyond predictable contractions to the 95% CIs). Reducing or increasing the number of sample temperatures marginally worsened or improved fits, respectively. Shifting the chosen temperatures did affect model fits; we elected to consistently use temperatures that could reasonably be selected in experimentally protocols and which provided good approximations of the underlying TPC. Code for the full randomization of these choices (number of sample points, number of measurement temperatures, choice of measurement temperature) is available in the online code repository, but further exploration of that randomization is beyond the scope of this project. Note that sampling at temperatures above $T_c$ results in consistent errors (as the equilibrium fitness TPC does not exist here) and sampling at temperatures that incur negative fitness causes errors for fitting some phenomenological TPC formulations which do not allow for negative values (no such formulations were used here).

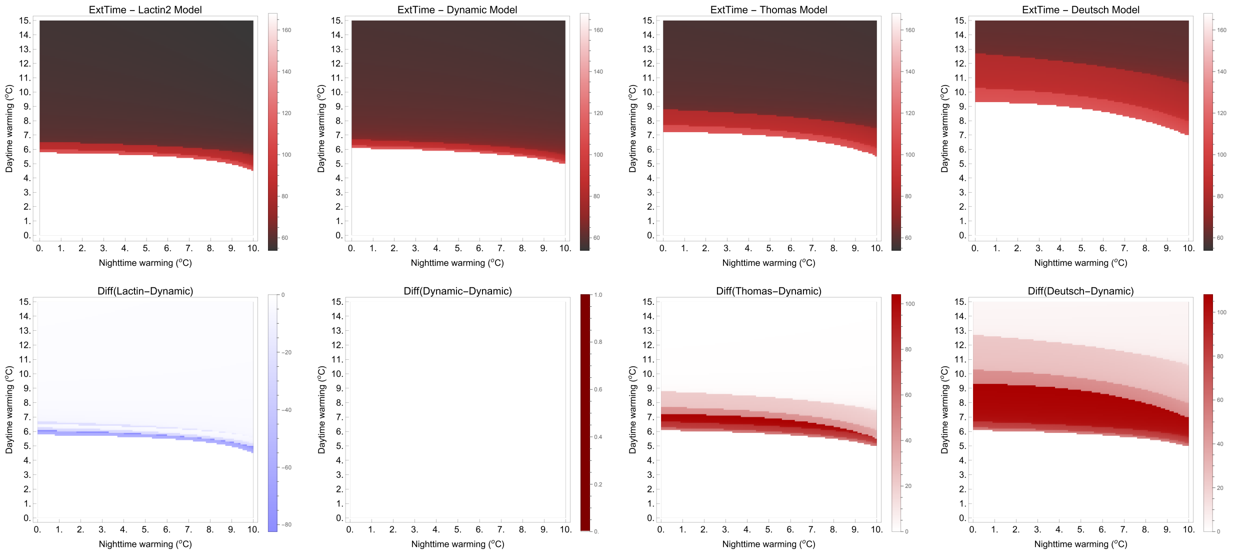


Figure S4: (a-d, top row) Projected extinction outcomes across the parameter space under different TPC model parameterizations for (a) the Lactin2 model, (b) the dynamic damage model, (c) the Thomas model, and (d) the Deutsch model. (e-h, bottom row) Difference plots comparing outcomes obtained using model fits compared to the dynamic model (calculated as phenomenological fit outcomes minus dynamic model outcome). Blue or red shading indicates an over- or under-prediction of risk, respectively, from the phenomenological model compared to the dynamic model.
